# Identifying Proteasome 26S Subunit, ATPase (*PSMC*) Family Genes as the Prognostic Indicators and Therapeutic Targets in Lung Adenocarcinoma

**DOI:** 10.1101/2022.04.23.489290

**Authors:** Md. Asad Ullah, Nafisa Nawal Islam, Abu Tayab Moin, Su Hyun Park, Bonglee Kim

## Abstract

This study explored the prognostic and therapeutic potentials of multiple Proteasome 26S Subunit, ATPase (*PSMC*) family of genes (*PSMC*1-5) in lung adenocarcinoma (LUAD) diagnosis and treatment. All the *PSMC*s were found to be differentially expressed (upregulated) at the mRNA and protein levels in LUAD tissues. The promoter and multiple coding regions of *PSMC*s were reported to be differentially and distinctly methylated, which may serve in the methylation-sensitive diagnosis of LUAD patients. Multiple somatic mutations (alteration frequency: 0.6-2%) were observed along the *PSMC*s coding regions in LUAD tissues that could assist in the high-throughput screening of LUAD patients. A significant association between *PSMC*s overexpression and LUAD patients’ poor overall and relapse-free survival (p<0.05, HR:>1.3) and individual cancer stages (p<0.001) was discovered, which justifies *PSMC*s as the ideal targets for LUAD diagnosis. Multiple immune cells and modulators (i.e., CD274, IDO1) were found to be associated with *PSMC*s expression in LUAD tissues that could aid in formulating *PSMC*-based diagnostic measures and therapeutic interventions for LUAD. Functional enrichment analysis of neighbor genes of *PSMC*s in LUAD tissues revealed different genes (i.e., *SLIRP, PSMA2, NUDSF3*) previously known to be involved in oncogenic processes and metastasis co-expressed with *PSMC*s, which could also be investigated further. Overall, this study recommends that *PSMCs* and their transcriptional and translational products are potential candidates for LUAD diagnostic and therapeutic measure discovery. However, further laboratory research is needed to validate the findings of this experiment.

## 1. Introduction

Lung adenocarcinoma (LUAD), which develops along the outer edge of the lungs within glandular cells in the small airways and falls under the umbrella of non-small cell lung cancer (NSCLC), is the most common type of histology, accounting for about 40% of all lung malignancies [1–2]. Worldwide research on 185 countries worldwide suggests that about 11.4% (more than 2.2 million) new cases of lung cancer were diagnosed in 2020, with an almost 18% mortality rate (1.8 million deaths) [3]. The low survival rate of patients with LUAD can be attributed to the lack of understanding of lung cancer biology, genomics and host factors that drive the progression of preinvasive lesions, heterogenicity of disease and patients’ outcomes, and so on. Although available diagnosis methods and treatment options have led to the overall decline in mortality rate from this prevalent cancer, the 5-years survival rate remains below 20% [4–5]. Investigation of specific prognostic biomarkers for disease stages or tumor types can help develop better screening strategies, improve patients’ prognoses, and assuage the financial burden of the disease [1,4,6–7]. Therefore, there is an increasing demand to secure an efficient diagnostic and therapeutic target for LUAD diagnosis and treatment that can significantly aid in the early-stage diagnosis, proper tracking of the patients throughout the cancer stages and appropriate therapeutic interventions ultimately reducing the medical burden.

The multiple Proteasome 26S Subunit, ATPase (*PSMC*) family of genes are reported to be involved in protein degradation, which plays a vital role in regulating the 26S proteasome [8]. This family of genes is comprised of six members, namely *PSMC1, PSMC2, PSMC3, PSMC4, PSMC5,* and *PSMC6* (*PSMC1-6*) [**Table 1**]. They partially constitute the formation of the 19S proteasome complex comprised of 19 essential subunits [9]. This regulatory complex, in turn, catalyzes the unfolding and translocation of substrates into the 20S proteasome [8].

**Table 1:** Genomic characteristics of all the subunits in PSMC gene family.

| Gene | Length of Nucleotide (bp) | Chromosomal Location | Length of amino acids | Protein encoded | Reference |
| --- | --- | --- | --- | --- | --- |
| *PSMC1* | 18,903 | 14q32.11 | 440 | 26S protease regulatory subunit 4 (26S proteasome AAA-ATPase subunit Rpt2) | 10,11 |
| *PSMC2* | 41,777 | 7q22.1-q22.3 | 433 | 26S protease regulatory subunit 7 (26S proteasome AAA-ATPase subunit Rpt1) | 10,12,13 |
| *PSMC3* | 7,705 | 11p11.2 | 439 | 26S protease regulatory subunit 6A (26S proteasome AAA-ATPase subunit Rpt5) | 10,14 |
| *PSMC4* | 10,615 | 19q13.11-q13.13 | 418 | 26S protease regulatory subunit 6B (26S proteasome AAA-ATPase subunit Rpt3) | 10,15,16 |
| *PSMC5* | 4,875 | 17q23.3 | 406 | 26S protease regulatory subunit 8 (26S proteasome AAA-ATPase subunit Rpt6) | 10,14,17 |
| *PSMC6* | 21,428 | 14q22.1 | 389 | 26S protease regulatory subunit S10B (26S proteasome AAA-ATPase subunit Rpt4) | 10,18,19 |

As of now, multiple *PSMC* family genes have been studied in the context of different human diseases including carcinoma. For example, previous study showed that *PSMC6* promotes osteoblast apoptosis and cancer cell proliferation by inhibiting the activation of the PI3K/AKT signaling pathway in an animal model of ovariectomy-induced osteoporosis [20]. *PSMC2* was found to be upregulated in osteosarcoma [21], prostate cancer [22], pancreatic cancer [23], glioma [24], oral squamous cell carcinoma (OSCC) [25], and hepatocellular carcinoma (HCC) [26]. Moreover, *PSMC2* was also reported to promote proliferation and inhibit apoptosis of glioma cells, and its knockdown halted the development and metastasis of prostate cancer [22] and progression of OSCC cells by promoting apoptosis via PI3K/Akt pathway and increasing the expression of pro-apoptotic proteins [25]. *PSMC5* is involved in the ubiquitination-dependent degradation of Tln1 and angiogenesis by blocking the miR-214/PTEN/Akt pathway [27]. Knockdown of Proteasome 26S subunit ATPase 3 interacting protein (*PSMC3IP*) resulted in the suppression of xenograft proliferation and tumorigenesis in the HCC cells [28]. In a recent study, researchers elucidated the crucial role of *PSMC* family members and their downstream-regulated genes in breast cancer progression [8]. However, the collective potential of the PSMC family of genes as candidates to be novel prognostic biomarkers and therapeutic targets in LUAD remains to be unveiled. Out of the six *PSMC* subunits, a recent study systematically evaluated the differential expression levels and prognostic values of *PSMC6* as a high *PSMC6* expression was associated with poor prognosis of LUAD, indicating the potential of *PSMC6* as a promising therapeutic target for LUAD [29].

This study sought to evaluate the prognostic and therapeutic significance of multiple Proteasome 26S Subunit, triple-A ATPase (*PSMC*) family of genes (*PSMC*1-5) in LUAD utilizing different web-based database mining approach (**Figure 1**). Since, the prognostic and therapeutic value of PSMC6 has been studied in the context of LUAD, this member of PSMC was not considered in this study [29]. Our study should contribute to understanding the predictive roles of *PSMCs* and their transcriptional and translational products in LUAD development, progression, and prognosis, which could be further validated using additional wet-laboratory experiments.

**Figure 1:**
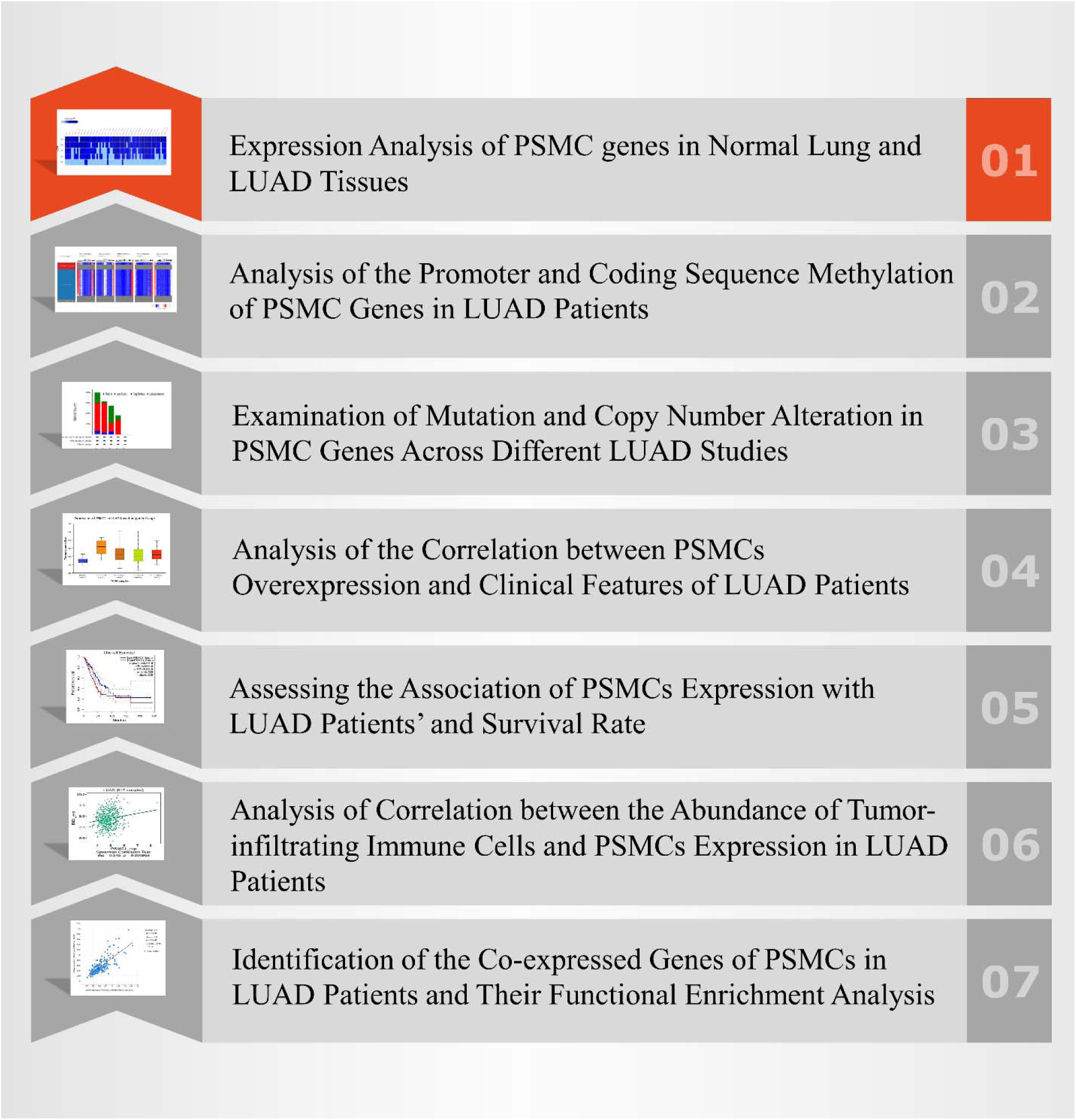
Strategies and tools utilized in the database mining approach employed in the overall study.

## 2. Materials and Methods

### 2.1. Expression Analysis of *PSMC* genes in Normal Lung and LUAD Tissues

*PSMC* gene expression pattern at the mRNA level in normal lung and cancerous LUAD tissues was determined using the OncoDB server (http://oncodb.org/, accessed on: April 7, 2022). OncoDB is an online platform that allows the users to explore the differential gene expression pattern in normal and corresponding cancerous tissues [1]. Thereafter, the Expression Atlas (https://www.ebi.ac.uk/gxa/home, accessed on: April 7, 2022) web-based tool was utilized to discover the mRNA level expression pattern of the *PSMC*s in a total of 68 different types of LUAD cell lines [2]. Finally, the Human Protein Atlas (HPA) (https://www.proteinatlas.org/, accessed on: April 7, 2022) server was used to determine the protein level expression of *PSMC*s in normal lung and LUAD tissues by analyzing the IHC images (at 200 µm length) of the post-mortem LUAD samples and adjacent normal lung tissues [3]. The Pathology and Tissues modules of the HPA server was explored to optimize the differences in PSMC protein expression in normal and cancerous lung tissues.

### 2.2. Analysis of the Promoter and Coding Sequence Methylation of *PSMC* Genes in LUAD Patients

The UALCAN server (http://ualcan.path.uab.edu/, accessed on: April 7, 2022) was utilized to examine the promoter methylation pattern of *PSMC* genes in LUAD tissues [4]. Integrated The Cancer Genome Atlas (TCGA) database was selected as the basis set for the experiment. Finally, the result was checked and validated based on a significant p value cutoff of <0.05. Next, the methylation pattern of the DNA sequence of *PSMC* coding genes was checked using the UCSC Xena browser (https://xenabrowser.net/, accessed on: April 7, 2022) [5]. In this step, again the integrated TCGA LUAD samples (n=706) were selected to observe the *PSMC* coding sequence methylation using the Methylation 450k array data. Finally, the GSCA server was used to confer the association between *PSMC* methylation and gene expression in LUAD tissues (http://bioinfo.life.hust.edu.cn/GSCA/#/, accessed on: April 7, 2022) [6]. The impact of *PSMC* methylation on the survival rate of LUAD patients was also evaluated from GSCA tool.

### 2.3. Examination of Mutation and Copy Number Alteration in *PSMC* Genes Across Different LUAD Studies

The cBioPortal server (https://www.cbioportal.org/, accessed on: April 7, 2022) was accessed to analyze the mutation and copy number alteration (CNA) in *PSMC* genes across a wide number LUAD study samples [7]. The data deposited by MSKCC, Broad, OncoSG, TDP, CPTAC and others including more than 2,598 patients’ samples over 9 overall studies were searched for *PSMC* mutation and CNA analysis in LUAD patients. The OncoPrint summary of the overall mutations of the selected *PSMC*s in different LUAD studies was inspected. Next, the bar diagram representing the type of genetic alterations in *PSMC* coding genes was also analyzed. Thereafter, the relation between the Overall Survival (OS) of LUAD patients and *PSMC* gene alteration was also evaluated from this server. Finally, the correlation between the CNAs present in *PSMC*s and mRNA expression in LUAD tissues was discovered in the form of bubble plot using the mutation module in GSCA (accessed on: April 7, 2022) server.

### 2.4. Analysis of the Correlation between *PSMC*s Overexpression and Clinical Features of LUAD Patients

The association between the *PSMC* gene overexpression and LUAD patients’ clinical features and demographic status i.e., age, individual cancer stages and nodal metastasis status was evaluated from the UALCAN server again (accessed on: April 7, 2022). UALCAN is a comprehensive user-friendly online tool that enables users to access omics data in cancer biomarker discovery and target validation. The TCGA LUAD samples were selected for the association analysis with our gene of interest in this study. The result of the analysis was considered significant based on the p-value cutoff of <0.05 found in student’s t test and the expression profile was represented as box plots wit transcript per million (TPM) reads unit.

### 2.5. Assessing the Association of *PSMC*s Expression with LUAD Patients’ and Survival Rate

The association between OS of LUAD patients and *PSMC* expression was established using the GEPIA 2 server (http://gepia2.cancer-pku.cn/, accessed on: March 7, 2022) [8]. GEPIA2 involves 9,736 tumors and 8,587 normal samples from GTEx and TCGA projects of RNA sequencing data, and this tool facilitates different transcriptional analysis, i.e., the analysis of correlation and differential expression across different normal and tumor tissues. The parameter values were kept at default during the analysis using the GEPIA2 server. Finally, the relation between *PSMC*s expression and LUAD patients’ relapse free survival (RFS) was also determined using the PrognoScan (http://dna00.bio.kyutech.ac.jp/PrognoScan/, accessed on: April 7, 2022) server [9]. The result of the experiment was then analyzed based on p-value and hazard ratio of LUAD patients on the basis of differential level of *PSMC* expression represented in the Kaplan-Meier plot of survival analysis.

### 2.6. Analysis of Correlation between the Abundance of Tumor-infiltrating Immune Cells and *PSMC*s Expression in LUAD Patients

The association of abundance of immune cells with *PSMC* expression in LUAD tissues was determined utilizing the immune module of the GSCA database (accessed on: April 7, 2022). GSCA is a highly inclusive database that helps in the analysis of different genomic association features and cancer patients’ clinical outcome across different forms of cancer. Moreover, it also aids in the analysis of correlation between different gene expression, gene mutation and the expression level of 24 different types of immune cells. Every selected *PSMC* gene was queried against the abundance of immune cells like B Cells, CD8+ T Cells, CD4+ T Cells, Macrophage, Neutrophil, Natural Killer (NK) Cells in LUAD tissues. Finally, the association between *PSMC* expression and abundance of different immunomodulators such as CD274 and IDO1 in LUAD tissues was determined from the TISIDB server (http://cis.hku.hk/TISIDB/, accessed on: April 7, 2022) [10]. The result of immune cell and modulator’s infiltration level was analyzed on the basis of p-value and correlation coefficient.

### 2.7. Identification of the Co-expressed Genes of *PSMC*s in LUAD Patients and Their Functional Enrichment Analysis

The co-expressed genes of *PSMC*s were identified using the TCGA LUAD database (Firehose, Legacy) from the cBioPortal server (accessed on: April 7, 2022). Thereafter, the top 300 positively co-expressed genes of each *PSMC* were selected based on p-value and correlation coefficient which were then used to identify the overlapping neighbor genes utilizing the InteractiVenn online tool (http://www.interactivenn.net/, accessed on: April 7, 2022) [11]. The overlapping neighbor genes of *PSMC*s in LUAD tissues were then used in gene ontology terms i.e., biological process (BP), molecular function (MF), cellular component (CC) and Kyoto Encyclopedia of Genes and Genomes (KEGG) pathway analysis from the Enrichr server (https://maayanlab.cloud/Enrichr/, accessed on: April 7, 2022) [12]. The result of the functional enrichment analysis was then presented using the ImageGP online and publicly available tool (http://www.ehbio.com/ImageGP/, April 7, 2022) [13].

## 3. Results

### 3.1. mRNA and Protein Level Differential Expression of *PSMC* Genes in Normal Lung and LUAD Tissues

The mRNA level expression of *PSMC*s in normal lung and LUAD tissues was analyzed from the OncoDB server. All the *PSMC* genes showed higher expression level in LUAD tissues compared to the normal lung tissues (**Figure 2**). Moreover, *PSMC*4 (fold change (FC): 0.80) showed the highest difference of expression among the test and control group followed by *PSMC*5 (FC: 0.49), *PSMC*2 (FC: 0.40), *PSMC*3 (FC: 0.30) and *PSMC*1 (FC: 0.21). Thereafter, the expression pattern of our genes of interest across 68 different LUAD cell lines was observed and *PSMC*3 was discovered to be overexpressed in most of the selected cell lines followed by *PSMC*4 and *PSMC*2 (**Figure 2f**). On the contrary, as in par with the previous result, *PSMC*1 showed the least overexpression in all the LUAD cell lines. Overall, all the selected *PSMC*s showed higher expression level in HCC461 and NCI-H1819 cell lines. Thereafter, the protein level expression pattern of the *PSMC*s in LUAD and their corresponding normal tissues was analyzed from the HPA server. *PSMC*2 showed medium staining against the administered antibody (HPA049621) in normal lung tissues whereas its higher staining evidence was recorded in the LUAD tissues (**Figure 3**). *PSMC*3 demonstrated low staining pattern in the normal lung tissues and medium staining in the LUAD tissues. Moreover, both *PSMC*4 and *PSMC*5 exhibited medium staining in the normal lung tissues whereas high level of staining was observed in the LUAD tissues.

**Figure 2:**
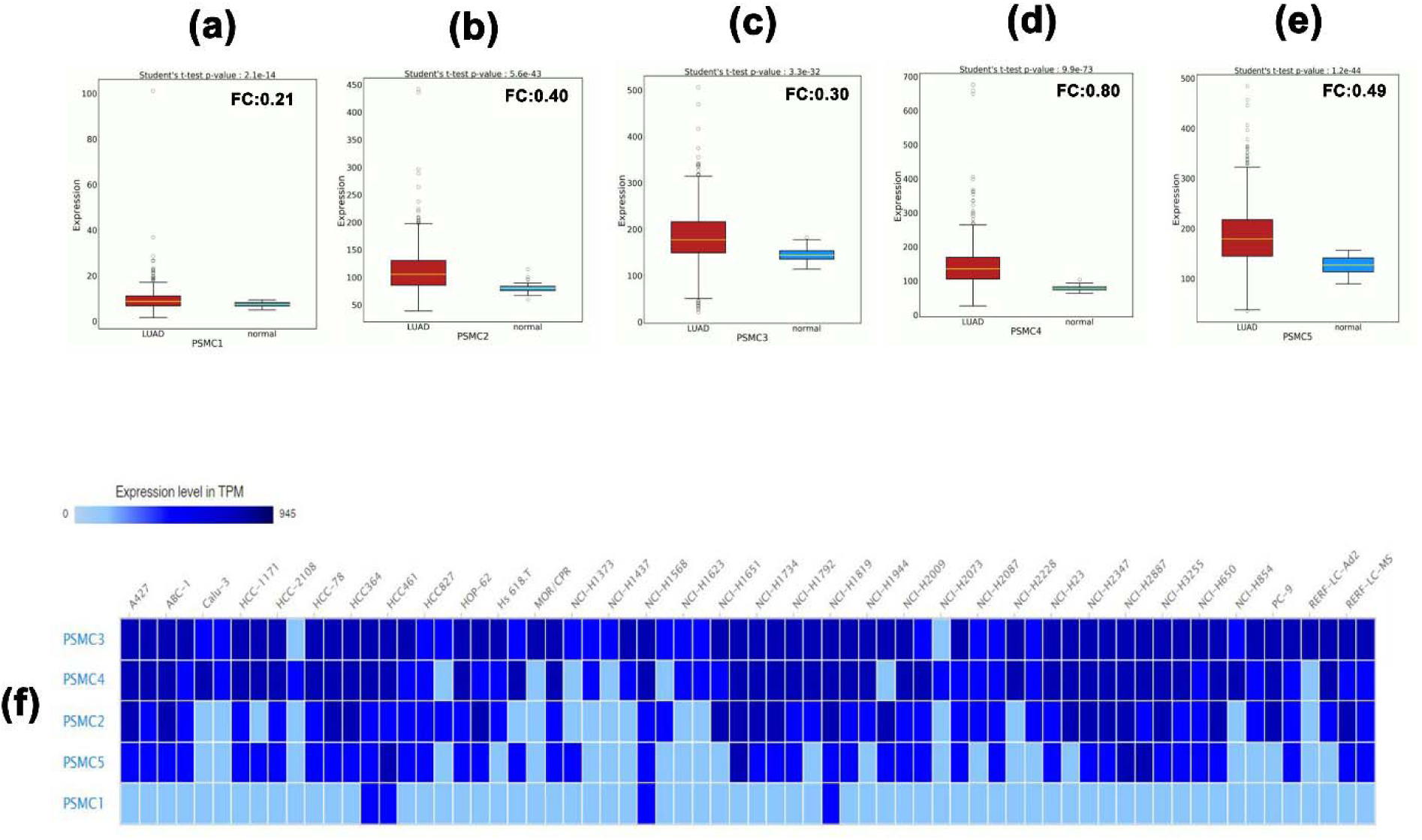
The mRNA level differential expression pattern of *PSMC*1 (a), *PSMC*2 (b), *PSMC*3 (c), *PSMC*4 (d) and *PSMC*5 (e) in normal lung and LUAD tissues observed from the OncoDB server. The expression of the *PSMC*s in different LUAD cell lines (f). FC: Fold Change.

**Figure 3:**
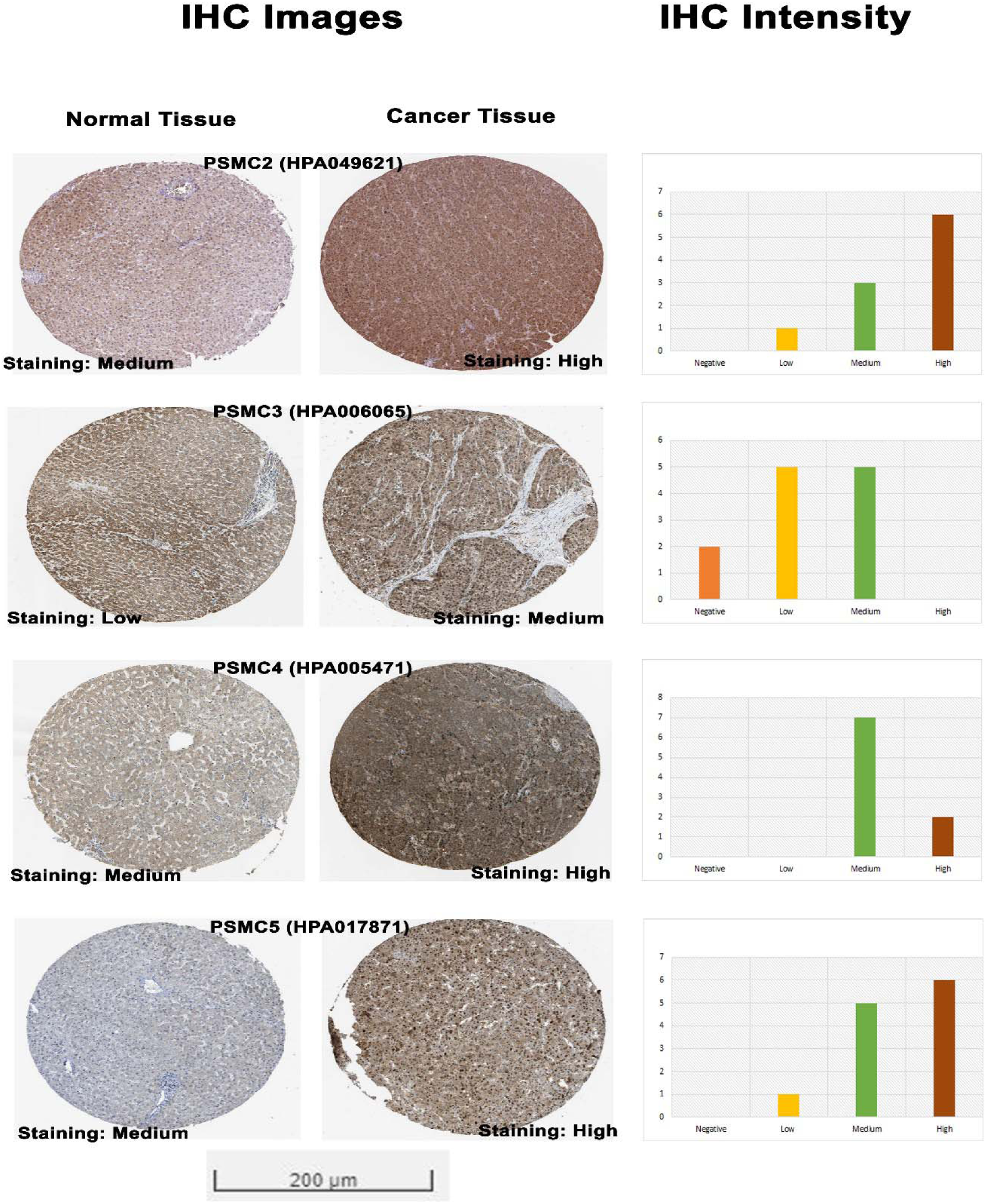
The IHC images (visualized at 200 µm) and IHC intensity delineating protein level expression of *PSMC*s in normal lung and LUAD tissue from the HPA server. The corresponding image for *PSMC*1 was not found.

### 3.2. Promoter and Coding Sequence Methylation Status of *PSMC* Coding Genes in LUAD Tissues

The promoter methylation pattern of the *PSMC* genes in LUAD and normal lung tissues was examined from the UALCAN server. *PSMC*1 gene coding promoter in LUAD tissues was found to be less methylated than in the normal lung tissues (p=3.76e-02) (**Figure 4**). Although, the *PSMC*2 and *PSMC*3 promoters were observed to be less methylated in LUAD tissues but the association was not significant (p>0.05). Additionally, *PSMC*4 (p=2.52e-09) and *PSMC*5 (p=6.50e-03) promoters were also found to be less methylated in LUAD tissues compared to the normal lung tissues. The coding sequence methylation analysis of the *PSMC*s in LUAD tissues from the UCSC Xena browser revealed that the selected *PSMC*s might have distinct coding sequence methylation pattern. For example, *PSMC*2, *PSMC*3 and *PSMC*4 signified that their coding regions might have the mostly methylated regions at the 3’ end of the sequence as indicated by the elevated beta value in red colored regions (**Supplementary Figure S1**). On the contrary, *PSMC*1 showed completely different pattern of methylation in which the most of the CpG islands might cover the 5’ end and a slight upstream region from the 3’ end of the coding sequence. In case of *PSMC*5 methylation pattern, the red landscapes at the 5’ end indicated that the initial region of the coding sequence might be hypermethylated. Finally, the effect of methylation in *PSMC* genes on their mRNA expression level specific to LUAD tissues was determined from the GSCA server. Unsurprisingly, methylation was negatively correlated with the *PSMC*1, *PSMC*2, *PSMC*4 and *PSMC*5 mRNA expression in LUAD tissues (**Supplementary Figure S2**). Additionally, the *PSMC*5 methylation was also found to be responsible for poor OS (p=0.043) and RFS (p=0.017) in LUAD patients (**Supplementary Figure S3**).

**Figure 4:**
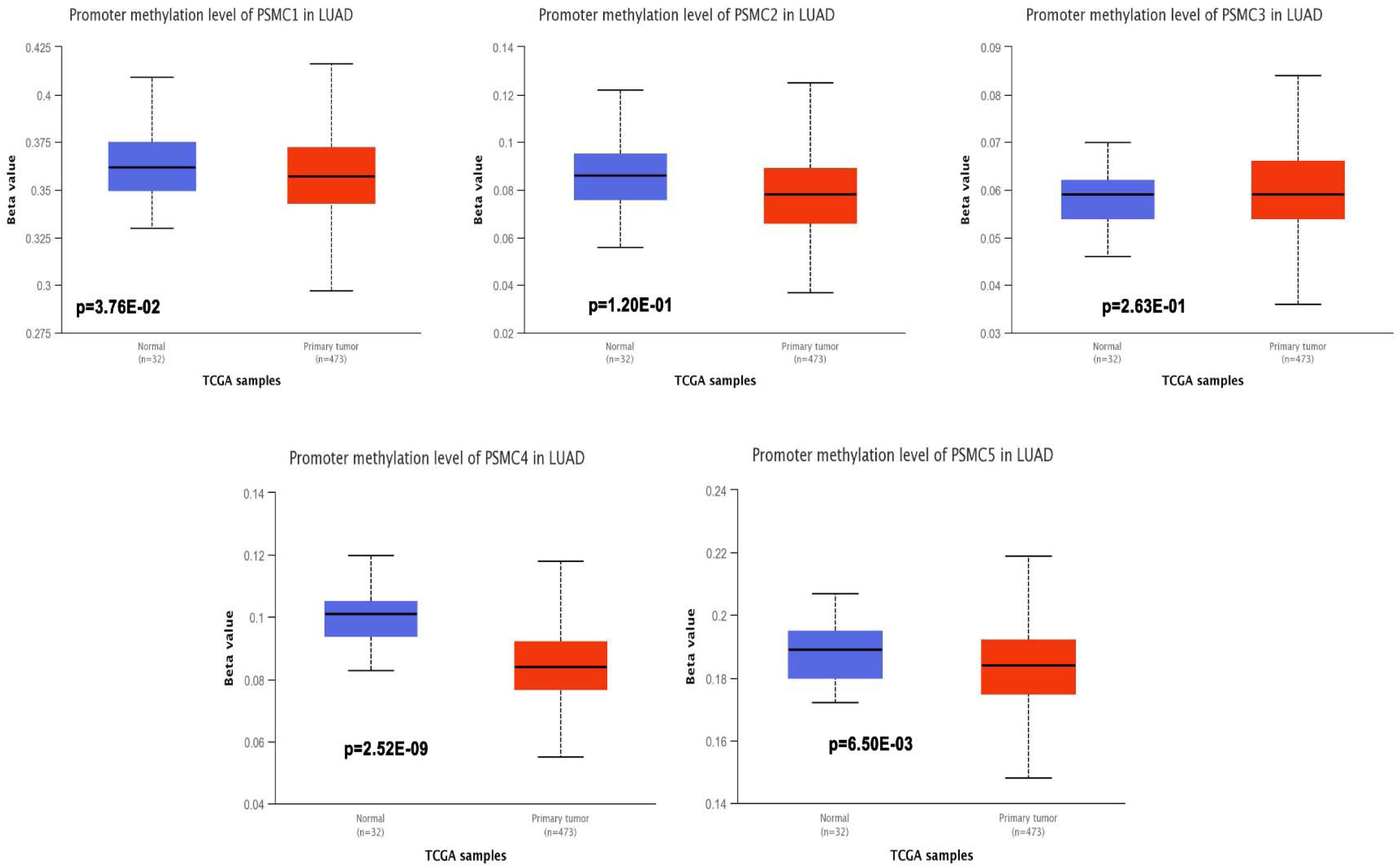
The promoter methylation pattern of *PSMC* genes in LUAD and normal lung tissues. Significant and distinct differential methylation pattern of different *PSMC*s in LUAD tissues was observed compared to the normal lung tissues.

### 3.3. Frequency of *PSMC* Mutation and Copy Number Alterations in LUAD Tissues

The mutation and copy number alteration frequency of *PSMC* genes across different LUAD studies was evaluated from the cBioPortal online tool. *PSMC*1, *PSMC*2, *PSMC*3, *PSMC*4 and *PSMC*5 showed an alteration frequency of 0.8%, 1.5%, 0.6%, 2% and 1.8%, respectively across different LUAD studies (**Figure 5a**). The analysis reported the presence of different detrimental genetic alterations i.e., amplification, deep deletion and splice sites across the *PSMC*coding regions in LUAD samples. Moreover, a number of missense mutations was also recorded in the *PSMC* coding regions carrying the potentials to interfere with the protein functions. The copy number alteration (CNA) frequency analysis revealed the evidences of a large number of amplification events across the selected LUAD studies responsible for the genetic alterations in *PSMC* genes in LUAD patients (**Figure 5b**). Thereafter, the association between *PSMC* CNA in LUAD tissues and their expression patterns was established from the GSCA server. All the *PSMC*s showed significant and positive correlation between the number of CNA events and their expression levels (**Figure 5c**). Finally, the effect of the mutations and CNAs present in *PSMC* genes on LUAD patients’ OS was also determined from this server. It was observed that *PSMC* mutations are significantly and negatively correlated to the LUAD patients’ OS (p=1.19e-04) (**Figure 5b**). Altogether, *PSMC*s altered LUAD patients had a poor OS (median survival ∼49 months) compared to the unaltered patients (median survival ∼66 months).

**Figure 5:**
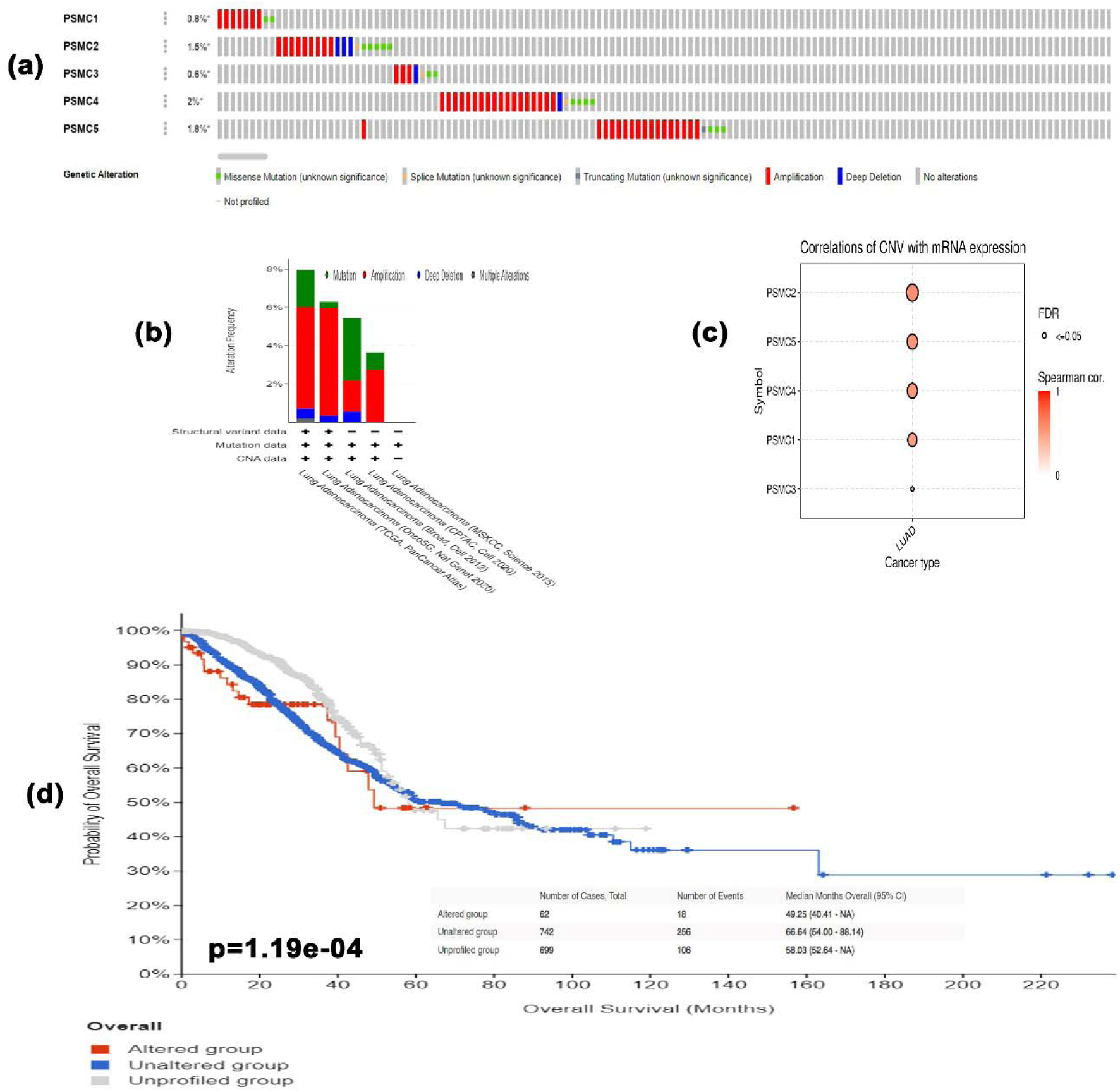
The mutation analysis report on *PSMC* genes in LUAD patients presented in OncoPrint diagram(a). The distribution of mutation and CNA in *PSMC* genes across different LUAD studies oriented in bar diagram (b). Ggplot representing the association between CNAs present in *PSMC* coding genes in LUAD tissues and *PSMC* expression levels. The association between *PSMC* mutation and LUAD patients’ OS represented in Kaplan-Meier plot (d).

### 3.4. Association between *PSMC*s Overexpression and LUAD Patients’ Clinical Features

The association between *PSMC*s overexpression and LUAD patients’ clinical characteristics was determined from the UALCAN server. Though, a noticeable rise on the *PSMC*s expression in the 21-40 years age group compared to the normal was found, *PSMC*1, *PSMC*3 and *PSMC*4 genes did not show significant association (p>0.05) (**Supplementary Table S1**) (**Figure 6**). Moreover, *PSMC*1-4 showed a marginal reduction in expression among other age groups except 21-40 years in LUAD tissues though the level remained above the normal lung tissue expression level whereas *PSMC*5 showed significant increase in expression in accordance with advancing age groups (p<0.05) (**Figure 6**). On the contrary, in terms of cancer stages, *PSMC*1 showed highest expression level in stage 1 and stage 4 whereas the intermediate stages showed a downward trend in expression in LUAD tissues although the expression level still remained above that in normal lung tissues (p<0.001) (**Supplementary Table S1**) (**Figure 6**). In case of other *PSMC*s, all the genes showed a marginal increment in their expression levels in comparison with aggressive cancer stages with a slight decline in the stage 4 in LUAD tissues (p<0.05). As in par with the *PSMC*1 expression level in comparison with cancer stages, its expression level was also found at the highest threshold in N0 and N3 of nodal metastasis status group in LUAD patients. However, the association between the expression level of *PSMC*1 and *PSMC*4 and N3 lymph node metastatic groups was not discovered to be significant (p>0.05) (**Supplementary Table S1**) (**Figure 6**). In contrast, other *PSMC*s sowed significant overexpression in accordance with the advancing metastasis stage in LUAD patients (p<0.04).

**Figure 6:**
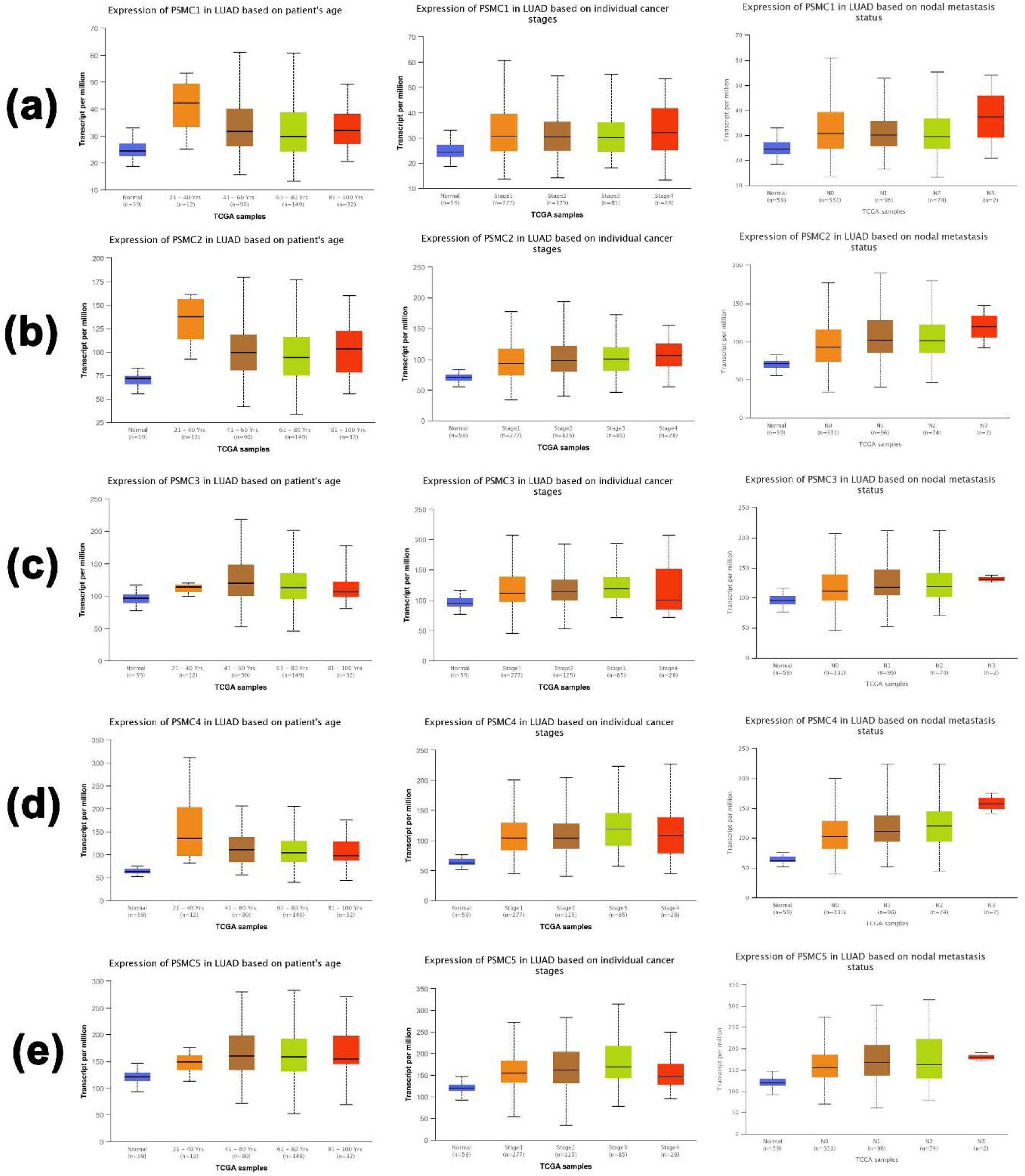
The pattern of *PSMC*1 (a), *PSMC*2 (b), *PSMC*3 (c), *PSMC*4 (d) and *PSMC*5 (e) overexpression in relation to LUAD patients’ age, individual cancer stages and nodal metastasis status represented in box plots. Several significant associations between *PSMC*s expression levels and LUAD patients’ clinical features was observed.

### 3.5. The Relation between *PSMC*s Expression and LUAD Patients’ Survival Rate

The relation between *PSMC*s expression and LUAD patients OS and RFS was established using the GEPIA 2 and PrognoScan servers, respectively. The report of the survival analysis was retrieved in the form of Kaplan-Meier plot. The analysis revealed that, *PSMC*1 overexpression is negatively correlated with the OS of LUAD patients (P=0.0016, Hazard Ratio (HR): 1.6) (**Figure 7a**) (**Supplementary Figure S4a**). Moreover, the *PSMC*1 overexpression was also responsible for the poor RFS of LUAD patients (P=0.03, HR: 3.0). Similarly, *PSMC*2 overexpression was discovered to be associated with the worsening OS (P=0.014, HR: 1.5) in patients (**Figure 7b**). Remarkably, *PSMC*2 overexpression was found to be the devastating cause of poor RFS in LUAD patients as observed by the heightened level of an HR of 13.49 (p=1.00e-16) (**Supplementary Figure S4b**). In case of *PSMC*3 expression, its overexpression was again significantly and negatively correlated to the poor OS (P=0.023, HR: 1.4) and RFS (P=1.00e-04, HR: 7.05) of the LUAD patients (**Figure 7c**) (**Supplementary Figure S4c**). Significant association was also observed for *PSMC*4 overexpression and the poor OS of LUAD patients from the report of the survival analysis (P=0.003, HR: 1.6) (**Figure 7d**). Again, the expression of *PSMC*4 was found to be attributed to the worse RFS in LUAD patients also (P=1.00e-05, HR: 4.95) (**Supplementary Figure S4d**). Finally, the *PSMC*5 expression was also discovered to be negatively associated with the OS (P=0.03, HR: 1.4) and RFS (P=0.015, HR: 7.73) of LUAD patients (**Figure 7e**) (**Supplementary Figure S4e**).

**Figure 7:**
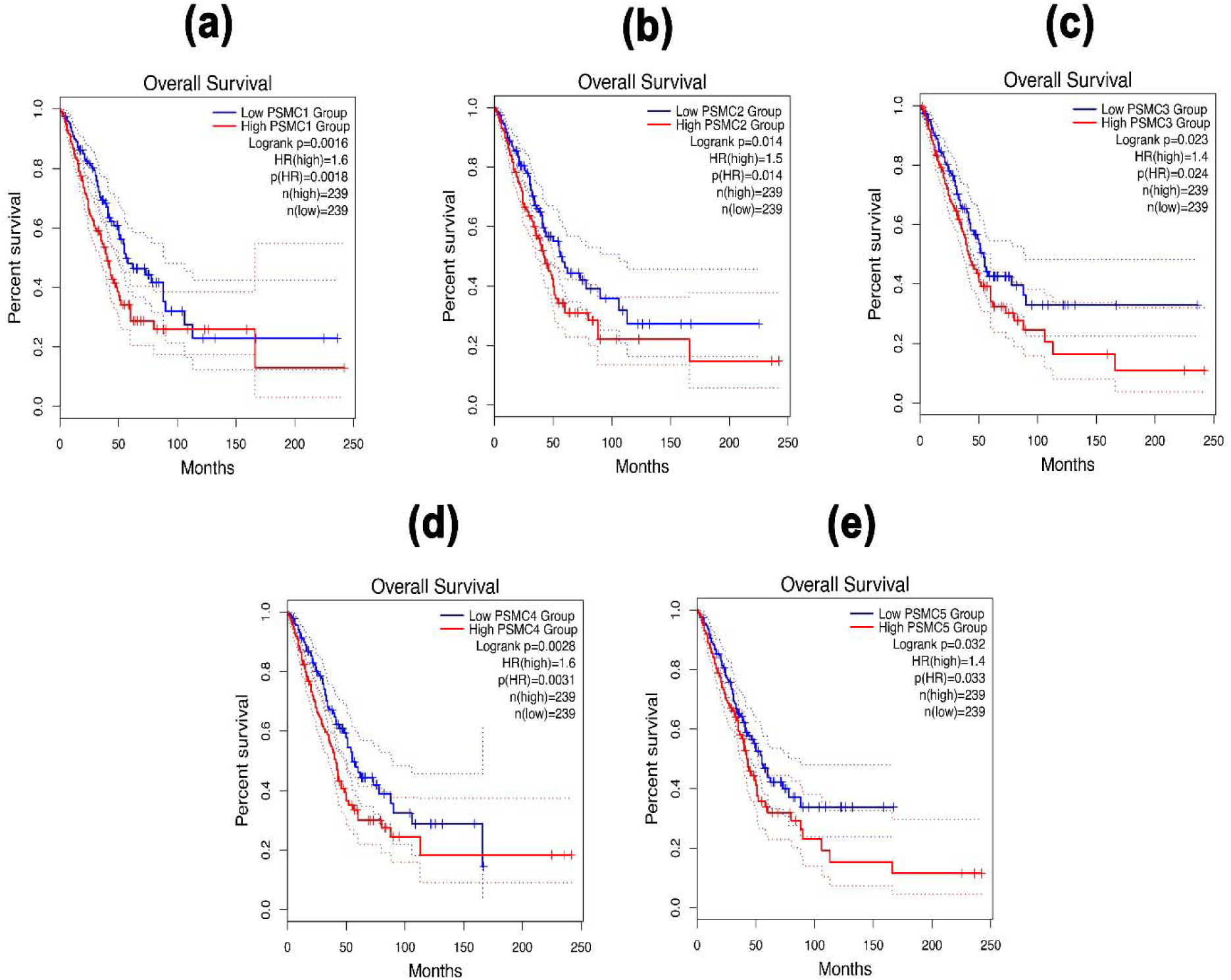
The Kaplan-Meier plot representation of *PSMC*1 (a), *PSMC*2 (b), *PSMC*3 (c), *PSMC*4 (d) and *PSMC*5 (e) expression and their relation with the OS of LUAD patients. Significant negative association was observed between *PSMC*s expression and LUAD patients’ OS (p<0.05, HR: <1.4).

### 3.6. The Association between *PSMC*s Expression and the Abundance of Tumor-infiltrating Immune Cells and Immune Modulators

The association between *PSMC*s expression and abundance of different immune cells was evaluated from the GSCA server. A significant positive correlation between *PSMC*1 expression and B cell (Cor: 0.09, p=0.016), CD8+ T cell (Cor: 0.13, p=0.009) was observed in LUAD tissues. However, negative correlation was observed between CD4+ T cell (Cor: -0.27, p=3.29e-11) and *PSMC*1 expression (**Supplementary Table S2**). *PSMC*2 expression showed significant association with the abundance level of B cell (Cor: 0.16, p=6.97e-05), CD8+ T cell (Cor: 0.09, p=0.02) and dendritic Cell (DC) (Cor: 0.21, p=1.13e-07). On the contrary, *PSMC*2 expression showed significant negative correlation with CD4+ T cell (Cor: -0.36, p=2.34e-19), Natural Killer (NK) cell (Cor: -0.14, p=0.0006) and Neutrophil (Cor: -0.12, p=0.02) abundance level in LUAD tissues (**Supplementary Table S2**). In case of *PSMC*3 expression, it showed significant positive association B cell (Cor: 0.14, p=0.004), CD8+ T cell (Cor: 0.15, p=0.002) and dendritic Cell (DC) (Cor: 0.12, p=0.003). However, *PSMC*3 exhibited negative association with CD4+ T cell (Cor: -0.28, p=7.01E-12), NK cell (Cor: -0.13, p=0.001) and Neutrophil (Cor: -0.08, p=0.04) infiltration levels in LUAD tissues (**Supplementary Table S2**). *PSMC*4 and *PSMC*5 showed positive association with B cell, CD8+ T cell, Monocyte and DC (Cor: > 0.09, p<0.05) abundance levels in LUAD tissues. A significant negative association with the CD4+ T cell, NK cell, Macrophage and Neutrophil production level and *PSMC*4 and *PSMC*5 expression levels was observed (p<0.05) **Supplementary Table S2**). Thereafter, the association between the *PSMC*s overexpression and immunoinhibitors like CD274 and IDO1 production level in LUAD tissues was established from the TISIDB server. *PSMC*1, *PSMC*2, *PSMC*4 and *PSMC*5 showed positive correlation with IDO1 expression levels in LUAD tissues (p<0.05) (**Figure 8a**). Moreover, the CD274 abundance levels in LUAD tissues showed negative association with *PSMC*1, *PSMC*3, *PSMC*4 and *PSMC*5 expression levels in LUAD tissues (p<0.05) (**Figure 8b**). Only *PSMC*2 showed positive association with CD274 infiltration levels.

**Figure 8:**
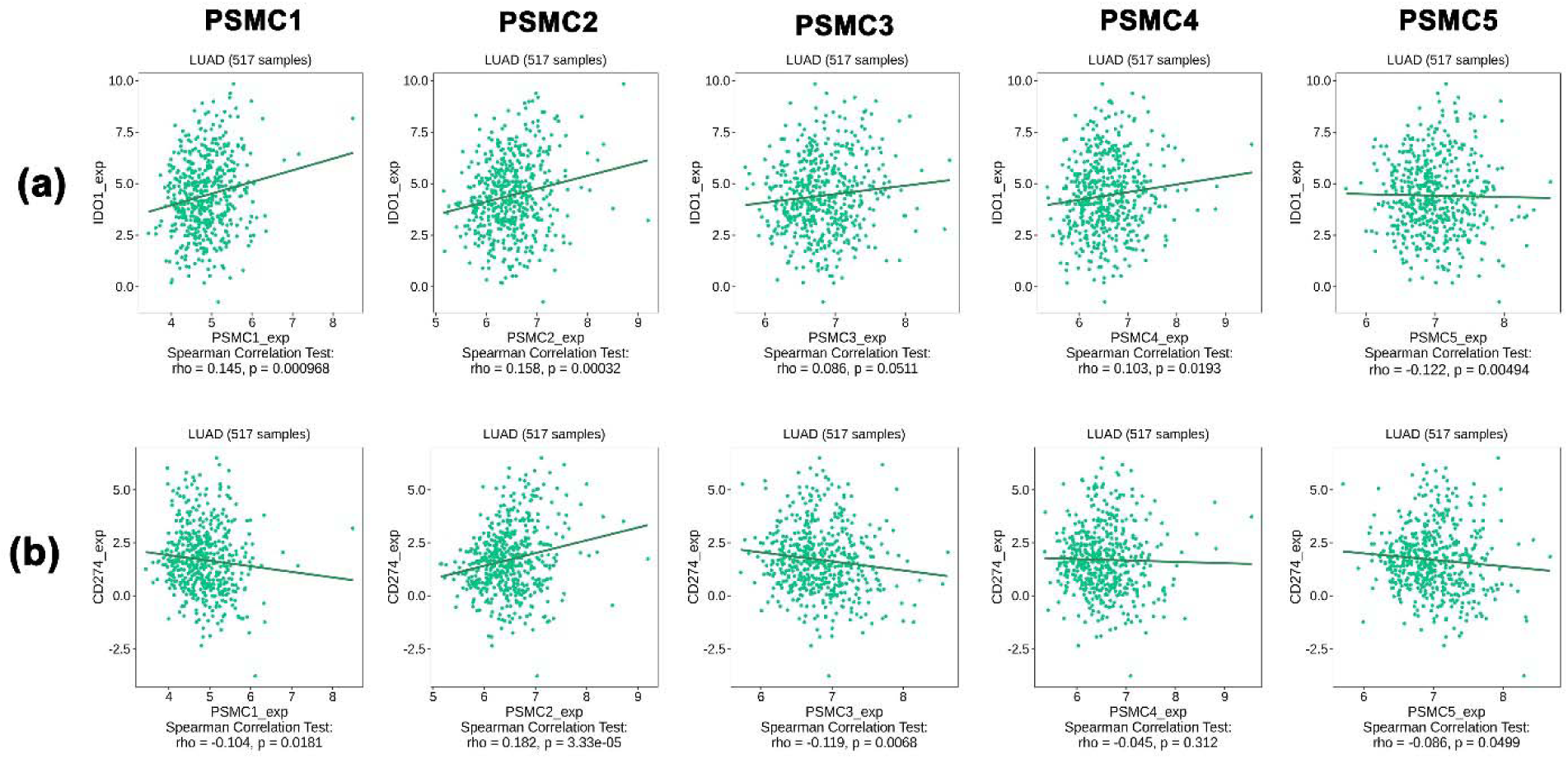
The association between *PSMC*s expression and the infiltration levels of IDO1 (a) and CD274 (b) in LUAD tissues. Significant positive and negative correlation was observed in between *PSMC* expression levels and immunomodulators expression levels.

### 3.7. The Co-expressed Genes of *PSMC*s in LUAD Tissues and Their Functional Enrichment Analysis

The top co-expressed gene of each *PSMC* in LUAD samples was identified from the cBioPortal server. *PSMC*1 was discovered to be highly co-expressed with SRA Stem-Loop Interacting RNA Binding Protein (*SLIRP*) coding gene in LUAD tissues (Cor: 0.76, p=4.30e-45) (**Figure 9a**). Proteasome 20S Subunit Alpha 2 (*PSMA2*) gene showed highest co-expression association with the *PSMC*2 gene (Cor: 0.76, p=4.30e-45) (**Figure 9b**). Moreover, *PSMC*3 gene was found to be most highly co-expressed with NADH: Ubiquinone Oxidoreductase Core Subunit S3 (*NDUFS3*) gene in LUAD tissue samples (Cor: 0.75, p=1.88e-42) (**Figure 9c**). Lastly, *PSMC*4 and *PSMC*5 genes were observed having highest level of co-expression with Translocase of Inner Mitochondrial Membrane 50 (*TIMM50*) (Cor: 0.75, p=5.99e-39) and Coiled-coil Domain Containing 137 (*CCDC137*) genes (Cor: 0.63, p=1.31e-26), respectively (**Figure 9d** and **9e**). Thereafter, the overlapping neighbor genes from top 300 positively co-expressed genes of each *PSMC* family member in LUAD tissues were identified using venn diagram (**Supplementary Figure S5**). The analysis revealed 13 genes i.e*., PSMB3, MRTO4, RFC2, TACO1, PRIM1, MCM3, KIF23, CCNA2, ERBB2, IRF1, PDCD45, SFN, TOX3* that are overlapped among the top 300 positively co-expressed genes of *PSMC*s. Afterward, the overlapping neighbor genes were investigated to understand their differential expression pattern in LUAD tissues. All the genes showed significant overexpression in LUAD tissues compared to the normal except *IRF1* that showed under-expression (**Supplementary Figure S6**). Finally, the overlapping genes were used in the functional enrichment analysis delineating different gene ontology terms i.e., biological processes, molecular functions and cellular component and KEGG pathway. The biological process analysis revealed that the highest ratio of the genes is involved in DNA replication, negative regulation of T cell differentiation, DNA metabolic processes, mitotic spindle assembly and so on (**Figure 10a**). The major molecular functions of the queried genes were DNA replication origin binding, phosphatase binding, motor activity and microtubule motor activity (**Figure 10b**). The overlapping genes were predominantly operating in intracellular membrane bound organelles, nucleus, and basolateral plasma membrane as observed from the cellular component analysis (**Figure 10c**). The KEGG pathway analysis on the overlapping neighbor genes of *PSMC*s in LUAD tissues reported that most of the genes are involved in pathways associated with bladder cancer, DNA replication, cell cycle, human papilloma virus infection and so forth (**Figure 10d**).

**Figure 9:**
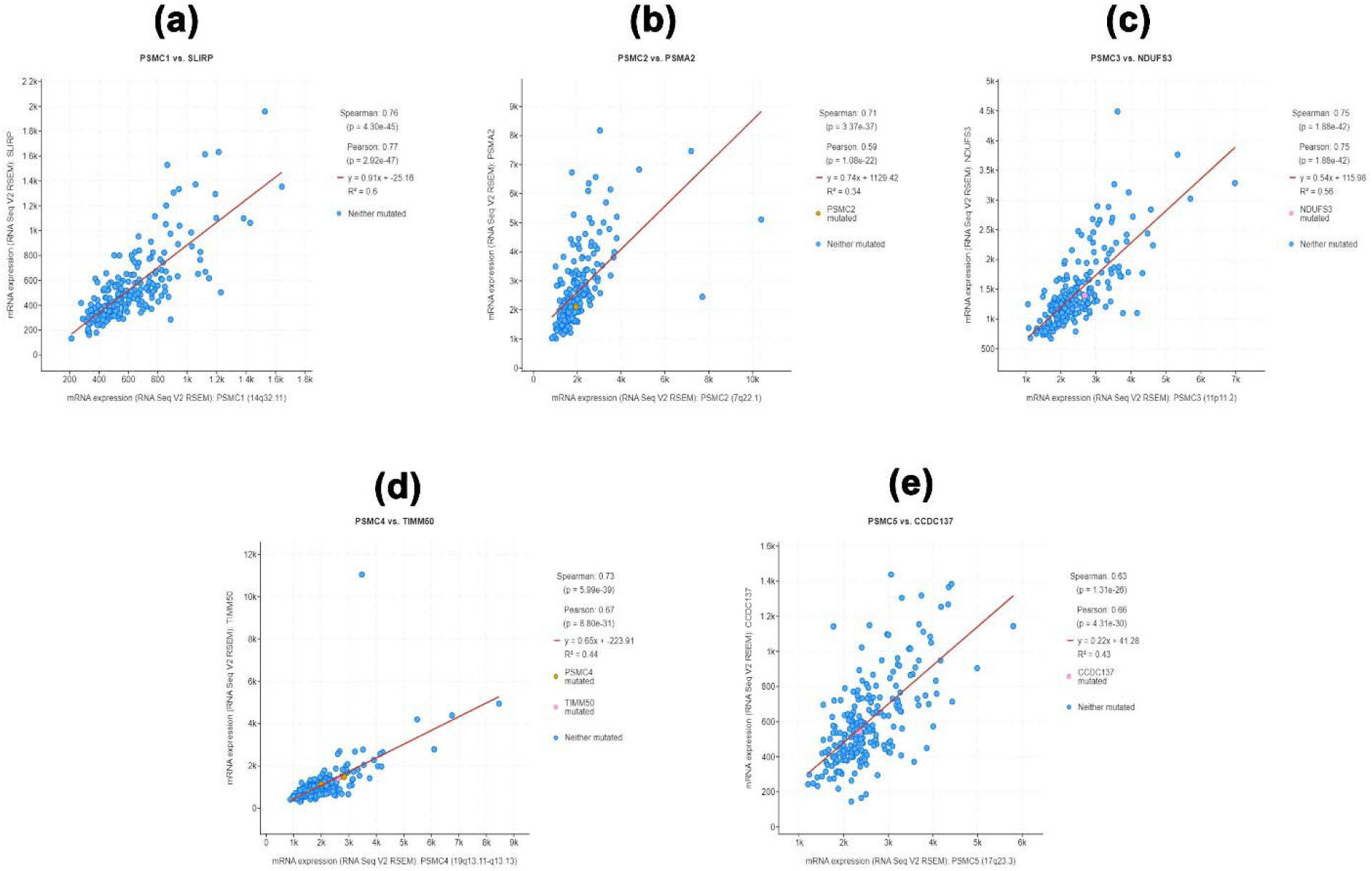
**T**he top positively co-expressed genes of *PSMC*1 (a), *PSMC*2 (b), *PSMC*3 (c), *PSMC*4 (d) and *PSMC*5 (e) in LUAD tissues.

**Figure 10:**
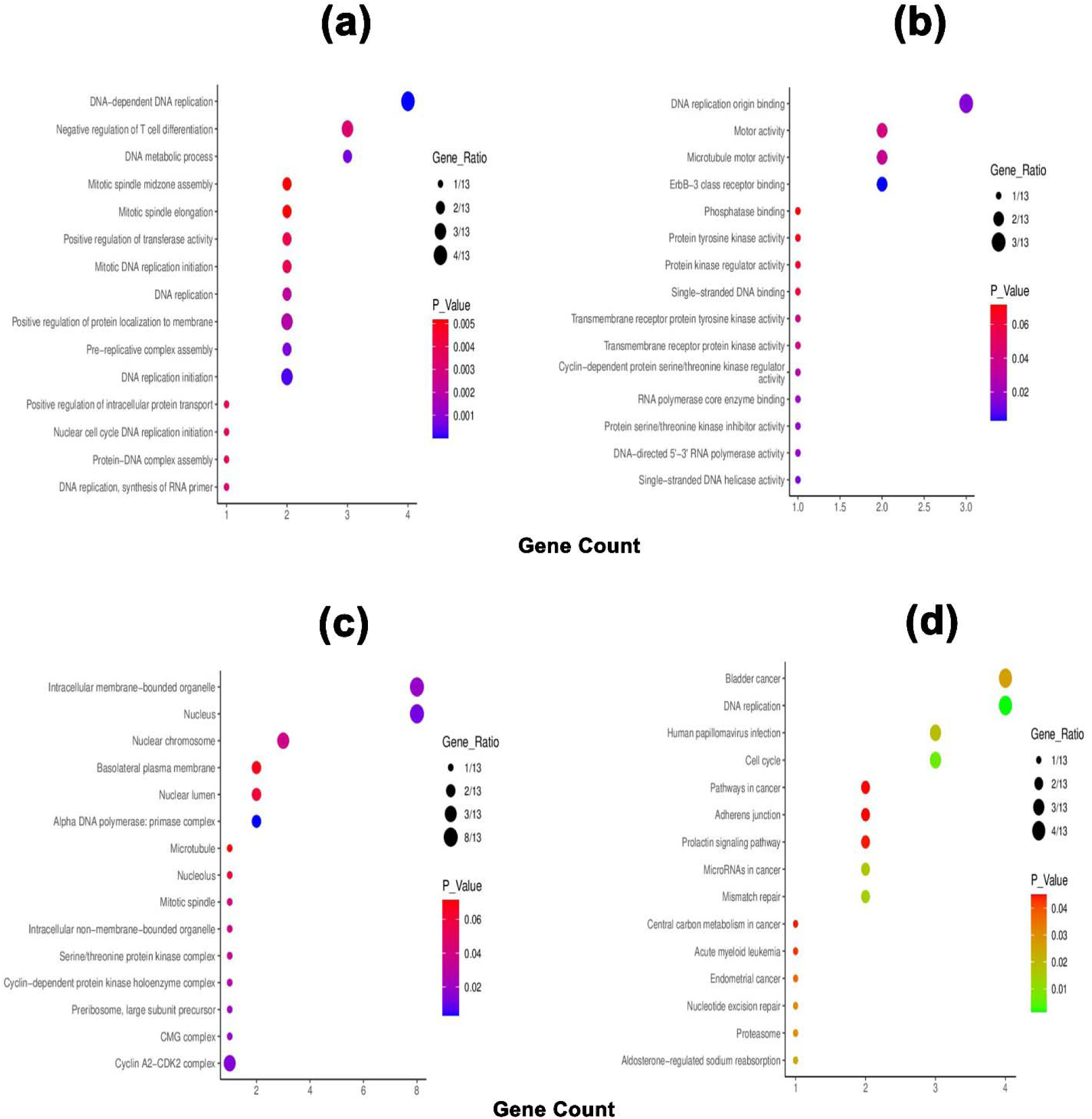
Bubble plots representing the top gene ontology terms of the overlapping neighbor genes of *PSMC*s in LUAD tissues: (a) biological processes, (b) molecular function, (c) cellular component and (d) KEGG pathway.

## 4. Discussion

This study explored the prognostic values of *PSMC* family of gene expression in LUAD taking the advantage of database mining approach. Initially, the differential expression pattern of all the selected *PSMC*s in LUAD and its corresponding adjacent normal lung tissues was evaluated. Given cancer development is a multistep process and is controlled by a variety of biological processes, differential gene expression analysis allows the understanding of the possible involvements of particular genes in the oncogenic development of a healthy cell [14–16]. In this study, we found that all the *PSMC*s are highly expressed in LUAD tissues compared to the normal lung tissues both at the mRNA and protein level suggesting their possible functions in LUAD development and progression (**Figure 2** and **3**). Moreover, all the genes of our interest also showed higher expression levels in different LUAD cell lines.

DNA methylation is one of the epigenetic drivers in cancer development and progression. Usually, promoter methylation regulates the gene activation and silencing and aberrant methylation is commonly associated with the up or downregulation of different genes in cancer cells [17]. Moreover, coding sequence methylation can also control the gene activity by altering the nucleosome orientation inside the chromatin structure [18, 19]. Additionally, the reduced methylation of different genes can propel the tumorigenesis of healthy cells inside the human body by escalating the activity of different oncogenes and thus differential methylation remains a promising target for epigenetic clinical decisions in cancer treatment [20]. In this experiment, the promoter regions of all the *PSMCs* were found to be differentially methylated (less methylated) in LUAD tissues compared to the normal lung tissues (**Figure 4**). Thus, the overexpression of the *PSMC*s in LUAD tissues may be attributed to the less methylated regions in *PSMC* coding promoters. In a recent study, *PSMC*5 methylation has been linked to being negatively associated with colorectal cancer exacerbation [21]. Furthermore, the evidence on the association between *PSMC*5 hypomethylation and LUAD patients’ poor OS and RFS reveals the potential of *PSMC*s in epigenetic-based therapeutic discovery for LUAD treatment (**Supplementary Figure S3**). Moreover, the differentially methylated circulating genes from samples like urine or blood of cancer patients can serve as a diagnostic marker for early-stage detection of lung cancer [22]. In this experiment, several different regions of *PSMC* coding sequences have been found to have distinct methylation patterns across LUAD samples which along with the differential level of promoter methylation may aid in the noninvasive diagnosis of LUAD patients (**Figure 4**).

Somatic driver mutations are the major etiological factors in the LUAD development and hence, the optimum understanding of the genetic alterations in relevant genes and their relations to patients’ survival is of paramount importance [23]. Unsurprisingly, CNA contributes more to the oncogenic development and subsequent growth of healthy cells than other nonsynonymous mutations like point mutations [24, 25]. In our study, all the *PSMCs* were predicted to have multiple somatic alteration events including amplification, deep deletion and splice which could promote the LUAD exacerbation by alternating the dosage of the translation products of the *PSMC*s inside the cells. In support of such assumptions, *PSMC* mutations in this study were associated with the poor OS of LUAD patients (**Figure 5**). What’s more, the presence of multiple missense mutations as evidenced in *PSMC* coding regions in LUAD patients may also influence the LUAD development and progression by producing non-functional, dysfunctional, or entirely no protein (**Figure 5**). Although the roles of *PSMC* gene mutations in cancer remain unstudied, multiple mutations in other proteasome family genes i.e., *PSMB5, PSMB6* and *PSMB7* are associated with the myeloma cell survival [26]. Apart from this, the prevalence of CNA events in different protein-coding genes can aid in the high-throughput diagnosis of lung cancer patients. For example, previously different CNAs including both loss and gain events in human chromosomes 3 and 6 have been successful in the high-throughput diagnosis of lung cancer within 44 months with an accuracy of 97% [27]. Thus, the observed alterations in *PSMC* genes in this study could also be investigated in formulating *PSMC*-based diagnostic measures for the early stage and accurate screening of LUAD patients.

Furthermore, all the *PSMC* genes were found to be overexpressed at an earlier age in the LUAD patients. A significant increase in the expression levels of the genes was observed across different cancer stages and with advancing lymph node metastasis status (**Figure 6**). The expression of *PSMC* genes was negatively associated with OS and RFS of LUAD patients (**Figure 7**). These pieces of evidence suggest that *PSMC*-based diagnostic measures may serve as an effective diagnosis method that could allow the early-stage diagnosis and tracking of LUAD patients throughout the clinical courses.

Tumor-infiltrating immune cells play a crucial role in inhibiting cancer cell growth and different immune cells have been shown to improve the prognosis of lung cancer patients [28]. Apart from this, the abundance of immune cells in cancer patients can also aid in tracking the patient’s status throughout the disease state and formulating immunotherapy for use during the clinical course [29]. For example, previous studies have shown that the abundance of CD4+, CD8+ T cells and Neutrophils are prognostic factors in lung cancer [30–32]. In this study, a significant association between *PSMCs* expression and different immune cells infiltration including the aforementioned ones was observed in LUAD patients that might assist in propagating dual diagnosis along with *the PSMC*-based diagnosis method (**Figure 7**). On the other hand, the mutated form of *PSMC* expression can alter the immune reactivity in the cancer microenvironment and worsen the prognosis of LUAD patients. Additionally, multiple *PSMC* mRNA expression levels were found to be positively and negatively correlated with the infiltration levels of different immunomodulators like CD274 (commonly known as programmed death-ligand 1; PD-L1) and indoleamine 2,3-dioxygenase 1 (IDO1) enzyme (**Figure 8**). PD-L1 is the most commonly found cell surface receptor in NSCLC and its overexpression predicts poor survival of lung cancer patients. Additionally, PD-L1 checkpoint inhibition and anti-PD-L1 antibodies are the most widely studied immunotherapy approaches in lung cancer, as well as, anti-PD-L1 antibodies are approved by Food and Drug Administration for IHC-based diagnosis of lung cancer [33, 34]. Moreover, IDO1 is a promising anticancer target for different cancer treatments including lung cancer whose function can be regulated by small candidate molecules and the process provides immune blockade opportunities outside the immune checkpoint inhibition and adoptive immune cell transfer [35].

The co-expression analysis revealed that *SLIRP* is the top highly co-expressed gene of *PSMC1* which was shown to have prognostic roles in colorectal cancer [36]. Among other selected top co-expressed genes, *PSMA2* was found to promote colorectal cancer cell proliferation and *NUDSF3* has been reported to be downregulated in the ovarian cancer cell and hypothesized to promote oncogenic development [37, 38] (**Figure 9**). *TIMM50*, a gene found to be highly co-expressed with *PSMC*4, promotes tumorigenesis and acts as a prognostic indicator in NSCLC [39]. Moreover, in a study involving 129 colorectal cancer (CRC) patients, *TIMM50* was discovered as a key regulator and prognostic marker of CRC [40]. Most recently, a pan-cancer analysis reported that *CCDC137* plays a crucial role and acts as a prognostic marker in different forms of cancers [41]. Given that the co-expressed genes are functionally related, all these shreds of evidence suggest that the *PSMC* family of genes might have an underlying mechanism in the oncogenic development of healthy lung cells.

Furthermore, 13 positively co-expressed overlapping genes of *PSMC*s in LUAD tissues were also found to be mostly associated with the LUAD development (**Supplementary Figure S6**). Gene ontology term analysis on these genes suggested that most of them are involved in DNA replication, controlling the cell cycle, and operating in the nucleus. The KEGG pathway analysis suggested that the genes are predominantly involved in different cancer pathways, development of bladder cancer, oncogenic virus infection pathways and so on. These findings again signify that the *PSMC* family genes may be associated with LUAD development and progression since the deregulation of the activity of *PSMC*s’ co-expressed genes in LUAD tissues can result in the initiation of oncogenic processes [42]. As a result, the co-expressed genes of the *PSMC*s could also be investigated in LUAD therapeutic and diagnostic measures discovery. However, further laboratory investigations are required on such assumptions.

In summary, this study demonstrated the differential expression of *PSMC*s in LUAD and showed variation in the methylation pattern of the *PSMC*s coding promoters and gene sequences between normal lung and LUAD tissues. A number of missense and truncating mutations were reported in the *PSMC*s coding regions. A significant association between *PSMC*s overexpression and LUAD patients’ clinical manifestation was observed. Moreover, *PSMC*s overexpression was correlated to the poor OS and RFS of the LUAD patients. These pieces of evidence suggest that *PSMC*s could be potential targets in the diagnosis and therapeutic interventions of LUAD patients. The association of *PSMC*s expression with different immune cells and immune modulators may aid in the further preparation of the *PSMC*-based diagnostic and therapeutic measures. The functional enrichment analysis unveiled different co-expressed genes controlling biological processes during the cell cycle. Thus, the neighbor genes of *PSMC*s could also be investigated further while extending laboratory work on making *PSMC*-based diagnostic and therapeutic measures for LUAD patients. Overall, this study suggests that *PSMC*s and their transcriptional and translational products are efficient prognostic and therapeutic targets for LUAD diagnosis and treatment. However, further laboratory research is warranted which is currently underway.

## Authors’ Contributions

Conceptualization and experimental designing: M.A.U. Experiment, analysis and interpretation: M.A.U., A.T.M. and N.N.I. Writing, review and editing: M.A.U., A.T.M., N.N.I., S.H.P. and B.K. Supervision and funding acquisition: B.K. All authors have read and agreed to the published version of the manuscript.

## Ethics Approval and Consent to Participate

Not Applicable

## Consent for Publication

## Availability of Data and Material

All the data are provided within the manuscript and the supplementary material.

## Competing Interest

All the authors declare that they have no conflict of interest regarding the publication of the paper.

## Funding

This research was supported by Basic Science Research Program through the National Research Foundation of Korea (NRF) funded by the Ministry of Education (NRF-2020R1I1A2066868), the National Research Foundation of Korea (NRF) grant funded by the Korea government (MSIT) (No. 2020R1A5A2019413), a grant of the Korea Health Technology R&D Project through the Korea Health Industry Development Institute (KHIDI), funded by the Ministry of Health & Welfare, Republic of Korea (grant number : HF20C0116), and a grant of the Korea Health Technology R&D Project through the Korea Health Industry Development Institute (KHIDI), funded by the Ministry of Health & Welfare, Republic of Korea (grant number : HF20C0038).

## Notes

### Competing Interest Statement

The authors have declared no competing interest.

