## Supplementary Information for "Identifying Proteasome 26S Subunit, ATPase (*PSMC*) Family Genes as the Prognostic Indicators and Therapeutic Targets in Lung Adenocarcinoma"

*Corresponding Author

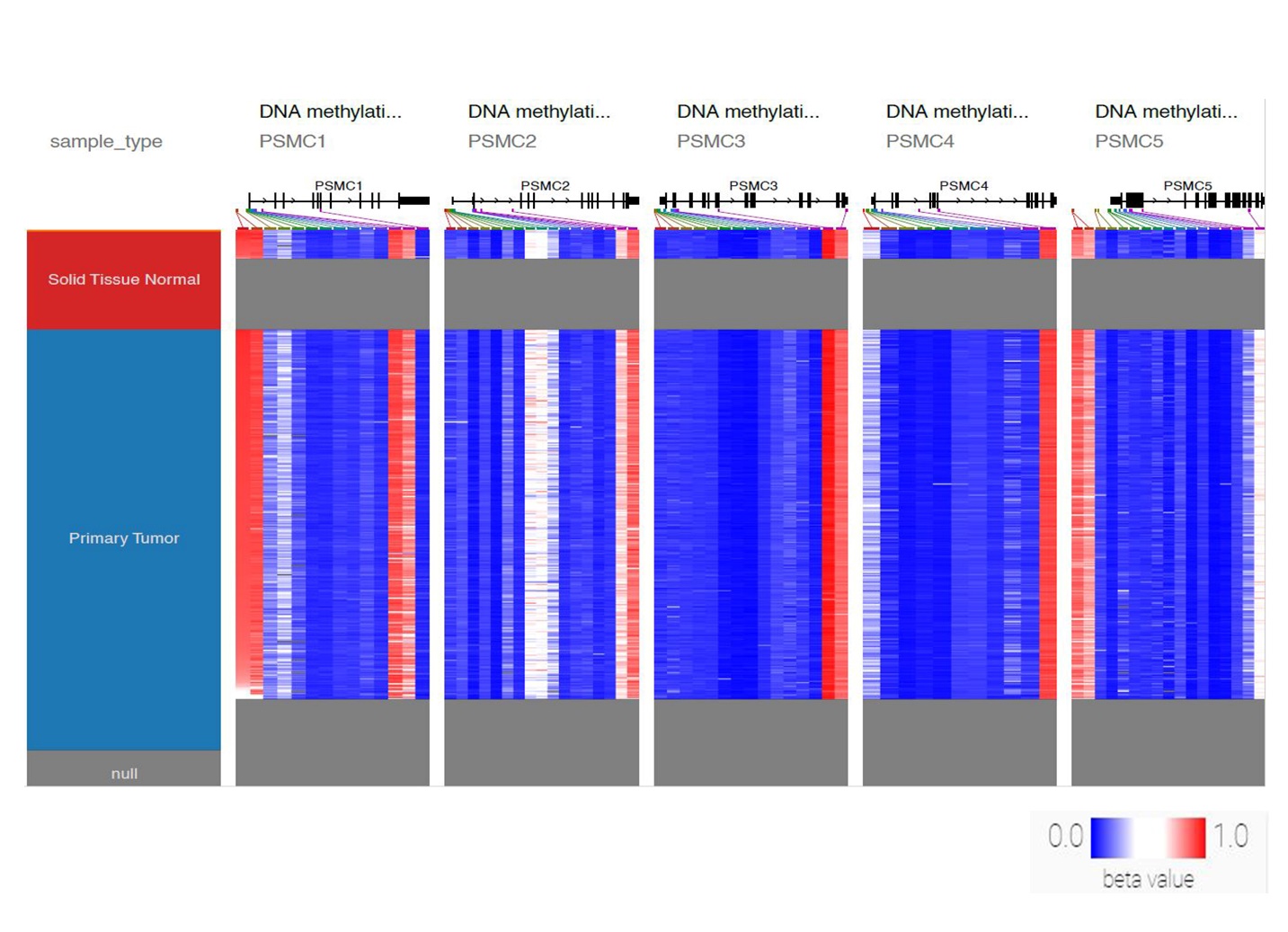
**Supplementary Information**

**Supplementary Figure S1:** The promoter coding sequence methylation pattern of PSMC genes in LUAD lung tissues in comparison with sample type. Significant and distinct differential methylation pattern of the PSMCs coding genes in LUAD tissues was observed. The grey region represents the lack of methylation data for the corresponding sample type in left-most bar.

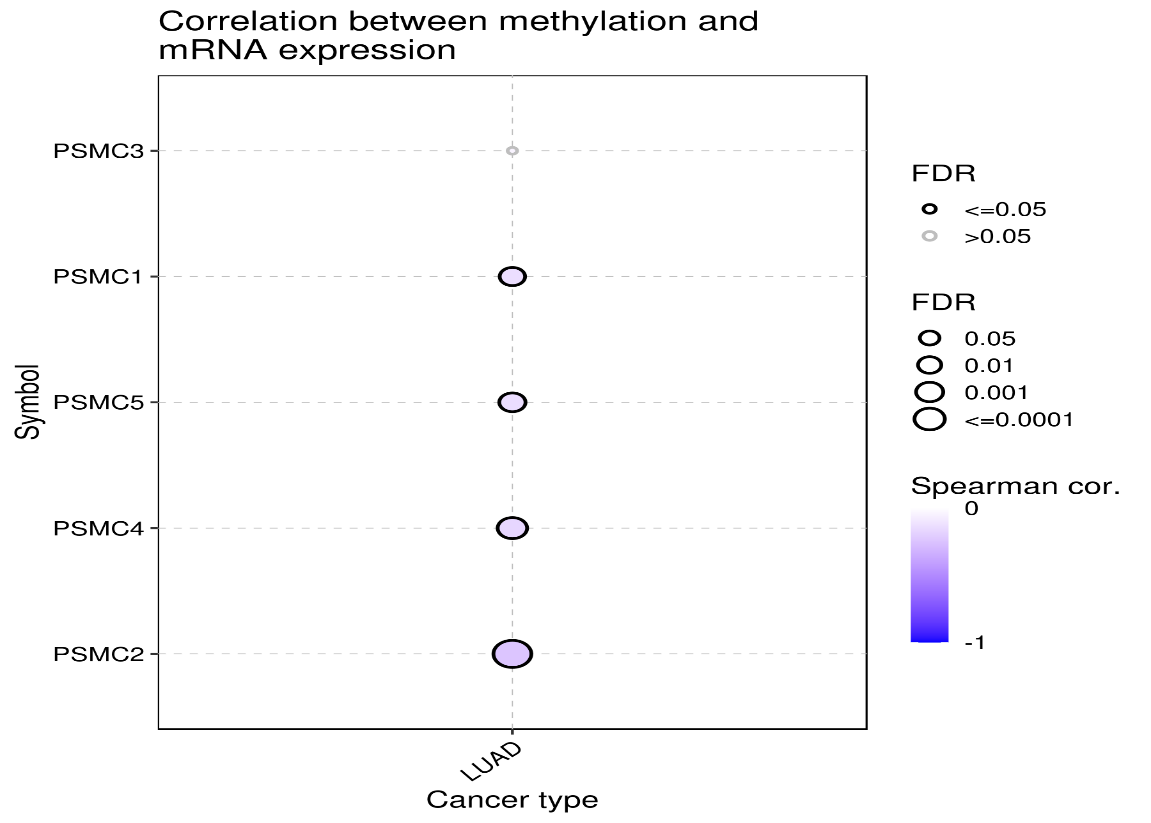

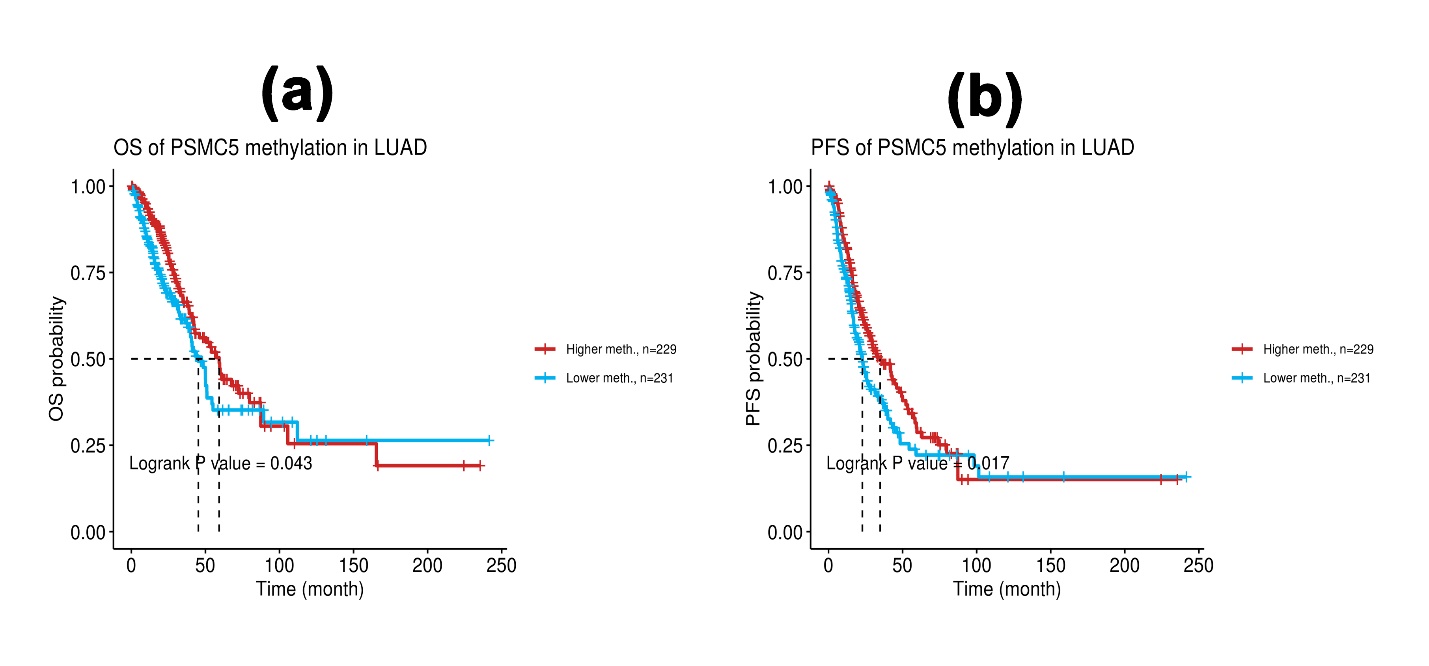
**Supplementary Figure S2:** The association between PSMCs methylation and mRNA expression in LUAD patients. Methylation level is negatively correlated with the mRNA expression level of PSMCs.

**Supplementary Figure S3:** The association between PSMCs methylation and LUAD patients’ OS and PFS. Lower methylation level of PSMC5 was negatively correlated to the OS and PFS of LUAD patients (p<0.05).

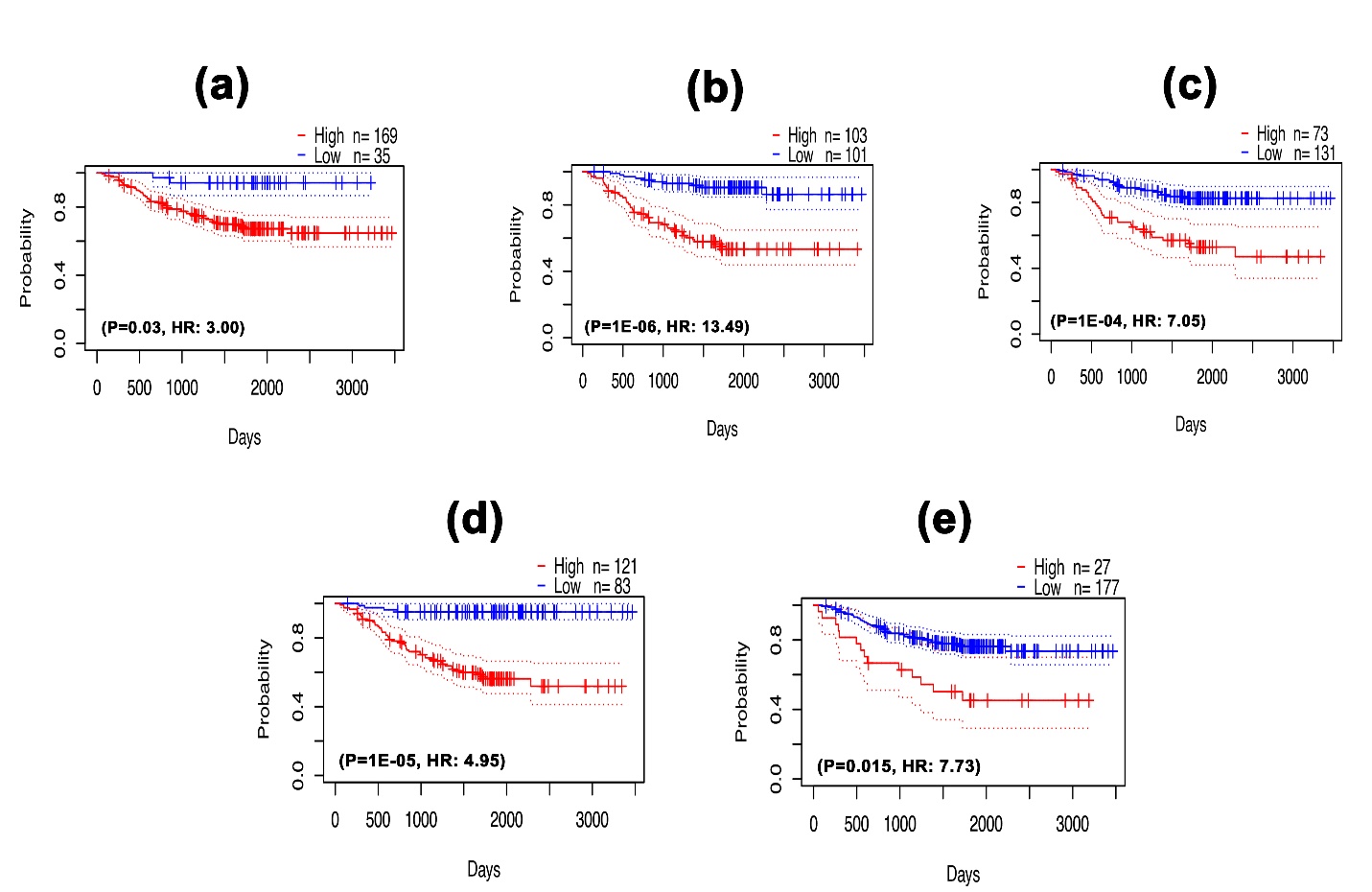

**Supplementary Figure S4:** The Kaplan-Meier plot representation of PSMC1 (a), PSMC2 (b), PSMC3 (c), PSMC4 (d) and PSMC5 (e) expression and their relation with the RFS of LUAD patients. Significant negative association was observed between PSMCs expression and LUAD patients’ OS (p<0.05, HR: >3.0).

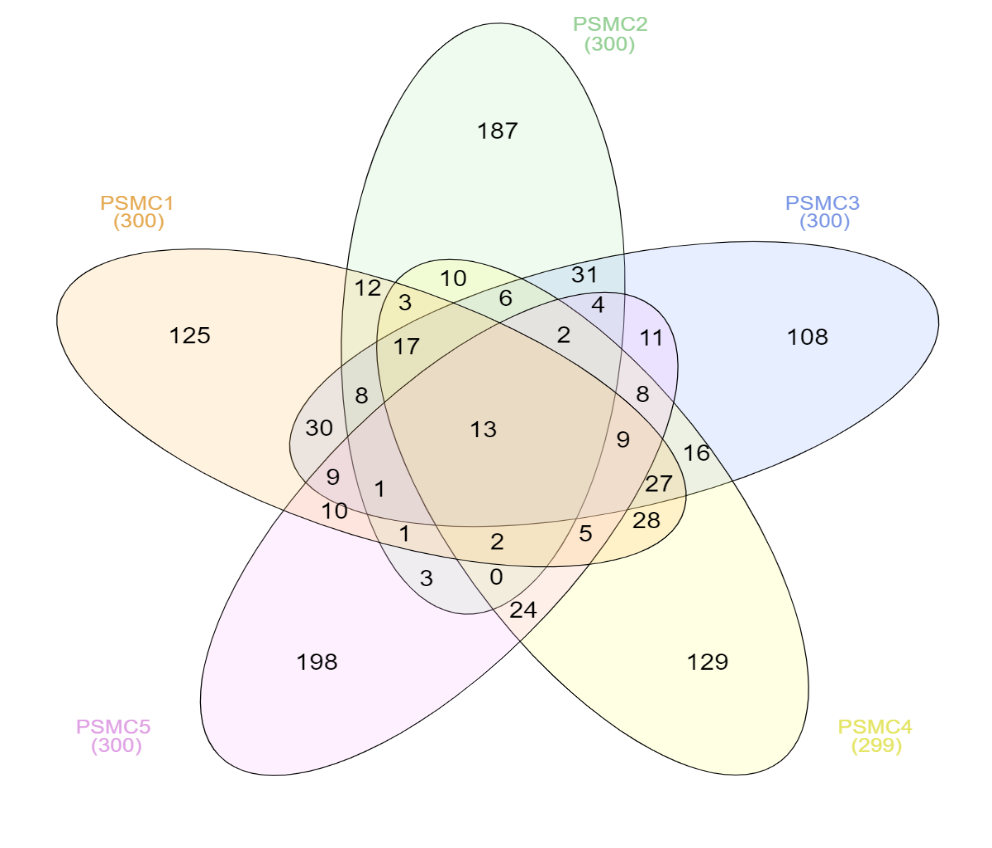

**Supplementary Figure S5:** Venn diagram representing the number of overlapping neighbor genes of PSMCs from top 300 selected co-expressed genes in LUAD tissues.

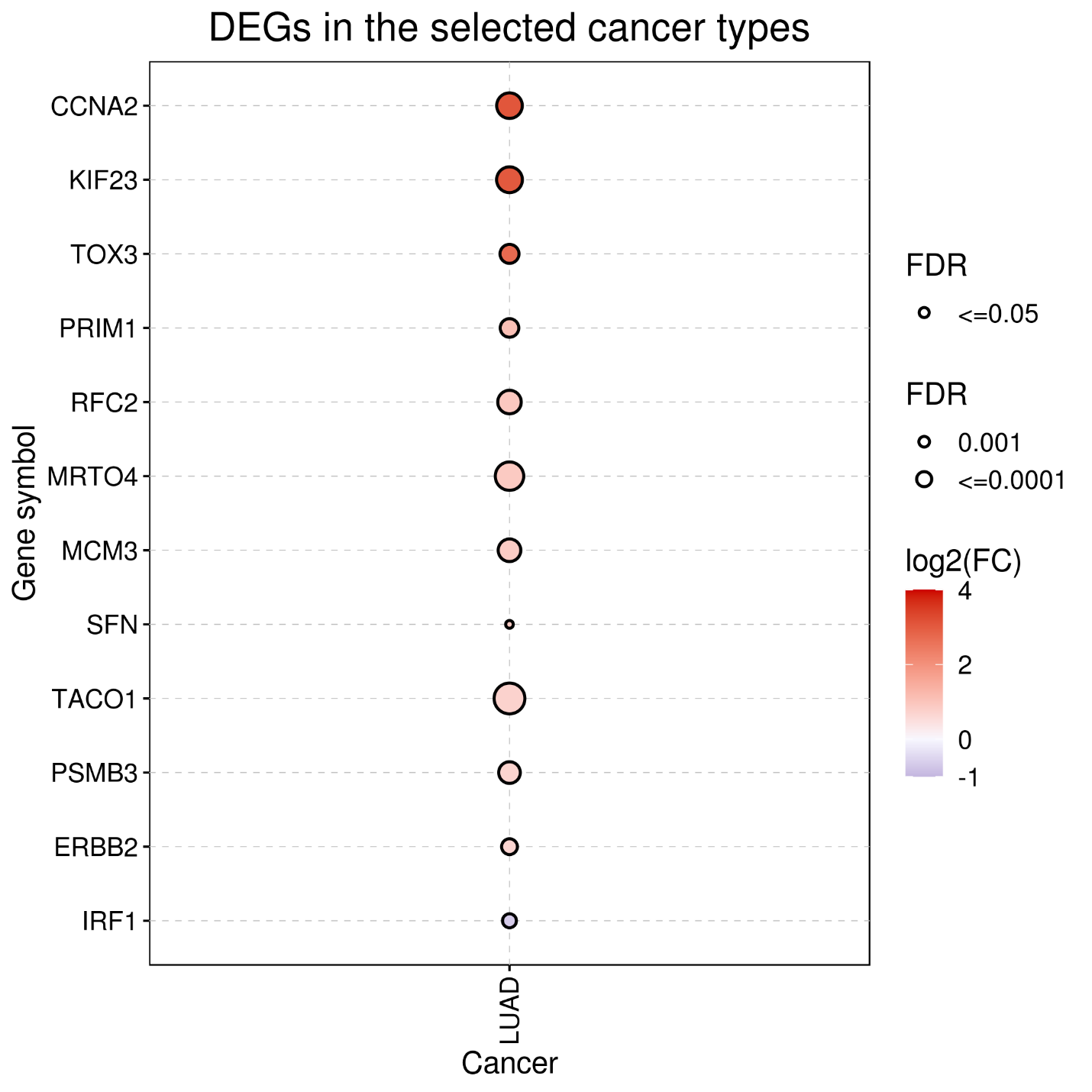

**Supplementary Figure S6:** The differential expression pattern of the 13 overlapping neighbor genes of PSMC in LUAD tissues. All the genes showed significant overexpression in LUAD tissues compared to the normal except IRF1 that showed under-expression.

| **Sl. No.** | **Features** | **Comparison** | **p-value** | | | | |
| --- | --- | --- | --- | --- | --- | --- | --- |
|  |  |  | ***PSMC1*** | ***PSMC2*** | ***PSMC3*** | ***PSMC4*** | ***PSMC5*** |
| 1. | **Age** | Normal-vs-Age(21-40Yrs) | **8.64E-02** | 3.21E-02 | **1.38E-01** | **1.45E-01** | 6.18E-04 |
|  |  | Normal-vs-Age(41-60Yrs) | 1.62E-12 | <1E-12 | <1E-12 | <1E-12 | <1E-12 |
|  |  | Normal-vs-Age(61-80Yrs) | 3.08E-10 | <1E-12 | <1E-12 | 1.62E-12 | <1E-12 |
|  |  | Normal-vs-Age(81-100Yrs) | 2.97E-04 | 5.64E-05 | 3.58E-03 | 1.29E-03 | 2.08E-05 |
| 2. | **Cancer Stages** | Normal-vs-Stage 1 | 1.62E-12 | <1E-12 | <1E-12 | <1E-12 | 1.62E-12 |
|  |  | Normal-vs-Stage 2 | 2.03E-08 | 3.04E-14 | 1.91E-12 | 1.62E-12 | 1.11E-16 |
|  |  | Normal-vs-Stage 3 | 7.71E-03 | 8.42E-11 | 3.17E-12 | <1E-12 | 1.24E-14 |
|  |  | Normal-vs-Stage 4 | 1.30E-03 | 1.73E-05 | 8.26E-03 | 3.54E-05 | 1.38E-04 |
| 3. | **Nodal Metastasis Status** | Normal-vs-N0 | 1.62E-12 | 1.62E-12 | 1.62E-12 | <1E-12 | 1.62E-12 |
|  |  | Normal-vs-N1 | 4.26E-07 | 1.62E-12 | 1.80E-12 | 1.62E-12 | 9.99E-16 |
|  |  | Normal-vs-N2 | 1.37E-02 | 2.96E-10 | 3.71E-10 | <1E-12 | 6.22E-12 |
|  |  | Normal-vs-N3 | **5.85E-01** | 3.35E-01 | 2.51E-06 | **1.15E-01** | 1.08E-07 |

**Supplementary Table S1:** The summary of the analysis revealing the association of PSMCs overexpression in relation to LUAD patients’ age, individual cancer stages and nodal metastasis status.

| **Gene Symbol** | **Immune Cell** | **Correlation Coefficient** | **P Value** | **FDR** |
| --- | --- | --- | --- | --- |
| ***PSMC1*** | B Cell | 0.099594 | 0.016802 | 0.03864 |
|  | CD4+ T Cell | -0.27175 | 3.29E-11 | 1.57E-10 |
|  | CD8+ T Cell | 0.137747 | 0.000918 | 0.003636 |
|  | DC | 0.079554 | 0.056368 | 0.111684 |
|  | Macrophage | -0.06857 | 0.100148 | 0.15146 |
|  | Monocyte | -0.01491 | 0.72105 | 0.816913 |
|  | NK | -0.07867 | 0.059176 | 0.091886 |
|  | Neutrophil | -0.04502 | 0.280715 | 0.420926 |
| ***PSMC2*** | B Cell | 0.164948 | 6.97E-05 | 0.000373 |
|  | CD4+ T Cell | -0.36278 | 2.34E-19 | 2.81E-18 |
|  | CD8+ T Cell | 0.094793 | 0.022893 | 0.054836 |
|  | DC | 0.218829 | 1.13E-07 | 1.3E-06 |
|  | Macrophage | 0.115703 | 0.005433 | 0.011418 |
|  | Monocyte | 0.062665 | 0.133054 | 0.242921 |
|  | NK | -0.14255 | 0.000601 | 0.001427 |
|  | Neutrophil | -0.12668 | 0.002318 | 0.00916 |
| ***PSMC3*** | B Cell | 0.146267 | 0.000429 | 0.001732 |
|  | CD4+ T Cell | -0.28058 | 7.01E-12 | 3.59E-11 |
|  | CD8+ T Cell | 0.152128 | 0.000248 | 0.001196 |
|  | DC | 0.12272 | 0.003177 | 0.010053 |
|  | Macrophage | -0.0156 | 0.708674 | 0.768672 |
|  | Monocyte | 0.035816 | 0.390894 | 0.53657 |
|  | NK | -0.13618 | 0.001051 | 0.00239 |
|  | Neutrophil | -0.08506 | 0.041277 | 0.097543 |
| ***PSMC4*** | B Cell | 0.216368 | 1.57E-07 | 2.09E-06 |
|  | CD4+ T Cell | -0.31289 | 1.51E-14 | 1.04E-13 |
|  | CD8+ T Cell | 0.208997 | 4.17E-07 | 4.64E-06 |
|  | DC | -0.03582 | 0.390802 | 0.516956 |
|  | Macrophage | -0.15234 | 0.000243 | 0.000678 |
|  | Monocyte | 0.095816 | 0.021455 | 0.057285 |
|  | NK | -0.16906 | 4.54E-05 | 0.000131 |
|  | Neutrophil | -0.11504 | 0.005708 | 0.019589 |
| ***PSMC5*** | B Cell | 0.125436 | 0.002563 | 0.007888 |
|  | CD4+ T Cell | -0.11052 | 0.007937 | 0.013264 |
|  | CD8+ T Cell | 0.081623 | 0.050233 | 0.104451 |
|  | DC | 0.030064 | 0.471442 | 0.594093 |
|  | Macrophage | -0.16563 | 6.49E-05 | 0.000203 |
|  | Monocyte | 0.113123 | 0.006572 | 0.021712 |
|  | NK | -0.16767 | 5.25E-05 | 0.00015 |
|  | Neutrophil | -0.0384 | 0.357639 | 0.501524 |

**Supplementary Table S2:** Summary of the association between PSMC mRNA expression different immune cell infiltration levels in LUAD tissues.
